# Classification of Single Particles from Human Cell Extract Reveals Distinct Structures

**DOI:** 10.1101/247254

**Authors:** Eric J. Verbeke, Anna L. Mallam, Kevin Drew, Edward M. Marcotte, David W. Taylor

## Abstract

Multi-protein complexes are necessary for nearly all cellular processes, and understanding their structure is required for elucidating their function. Current high-resolution strategies in structural biology are effective, but lag behind other fields (e.g. genomics and proteomics) due to their reliance on purified samples rather than characterizing heterogeneous mixtures. Here, we present a method combining single particle analysis by electron microscopy with protein identification by mass spectrometry to structurally characterize macromolecular complexes from extracts of human cells. We obtain three-dimensional structures of native proteasomes directly from *ab initio* classification of a heterogeneous mixture of protein complexes. In addition, we find an ~1 MDa size structure of unknown composition and reference our proteomics data to suggest possible identities. Our study shows the power of using a shotgun approach to electron microscopy (*shotgun EM*) when coupled with mass spectrometry as a tool to uncover the structures of macromolecular machines in parallel.

## Introduction

Protein complexes play an integral role in all cellular processes. Understanding the structural architecture of these complexes allows direct investigation of how proteins interact within macromolecular machines and perform their function. In an effort to understand why proteins form these machines, proteome-wide studies have been conducted to determine the composition of protein complexes (Drew et al., 2017a; Gavin et al., 2002; Havugimana et al., 2012; Hein et al., 2015; Ho et al., 2002; Huttlin et al., 2015, 2017; Kastritis et al., 2017; Kristensen et al., 2012; Krogan et al., 2006; Wan et al., 2015). Similar studies have identified direct contacts between protein complex subunits computationally (Drew et al., 2017b) or by cross-linking mass spectrometry (Leitner et al., 2016; Liu and Heck, 2015; Rappsilber et al., 2000), and while these studies provide insightful predictions on protein-protein interactions, they lack directly observable structural information that can inform us on function and subunit stoichiometry.

Structural genomics approaches, such as the Protein Structure Initiative, have thus far been the most successful way to systematically solve structures for proteins lacking a model (Chandonia and Brenner, 2006). These approaches have removed several bottleneck steps in traditional structural biology by applying high-throughput technology to sample preparation, data collection and structure determination. Many high-resolution structures have resulted from structural genomics. However, these approaches typically miss large complexes and perform best on single proteins or low molecular weight complexes that can be purified and crystallized for X-ray crystallography, or labeled for nuclear magnetic resonance (Montelione, 2012).

Recent advances in electron microscopy (EM) software and hardware have dramatically increased our ability to solve the structures of native protein complexes and allow for increased throughput approaches using EM. Automated microscopy software such as Leginon (Suloway et al., 2005), SerialEM (Mastronarde, 2005), and EPU (FEI) allow for the collection of large datasets in a high-throughput, semi-supervised manner. RELION, a Bayesian algorithm for 3D classification, allows users to sift through conformationally heterogeneous samples to define structurally homogenous classes (Scheres, 2012). Furthermore, 3D reconstructions can now be done *ab initio* (without an initial model) by a computationally unsupervised approach using cryoSPARC (Punjani et al., 2017). These strategies potentially allow for analysis of heterogeneous mixtures, although this aspect has not been explored extensively.

Advances in hardware, such as direct electron detectors and Volta phase plates, allow visualization of particles at near atomic resolutions and smaller molecular weights, which was previously only possible for larger particles or particles with high symmetry (Danev and Baumeister, 2016; Kühlbrandt, 2014). Despite these revolutionary advances, single particle EM is still largely used to study homogenous samples, where the identity of the protein complex is known *a priori*.

Here, we take a different approach to structure determination by exploiting advances in EM software to structurally classify native protein complexes from human cell lysate. By using a shotgun approach to EM (*shotgun EM*), we chromatographically separate cell lysate into tractable fractions before identification by mass spectrometry (MS) and structural analysis by EM. Using this approach, we characterize compositionally and structurally heterogeneous protein complexes from immortalized (HEK293T) cells separated by macromolecular size using size-exclusion fast protein liquid chromatography (SEC-FPLC).

The protein composition of a single, high molecular weight, sample was determined by running liquid chromatography tandem mass spectrometry (LC-MS/MS) experiments. Identified proteins were then mapped to previously generated protein interaction networks to reveal candidate protein complexes. We then collected negative stain EM data and performed single particle analysis of heterogeneous particles simultaneously. This approach reveals the structural information available from investigating cell lysate as opposed to a purified sample. Using this approach, we are able to identify the dominant and structurally distinctive macromolecular machines after unbiased 3D classification or *ab initio* reconstructions of single particles.

## Results

### Separation and identification of subunits from high molecular weight protein complexes

Intact complexes from lysed human cells were first separated by macromolecular size using SEC-FPLC. We selected a high molecular weight fraction (fraction 4) for MS and EM analysis (Figure 1) with molecular weights in the range of 1.5 to 2 MDa based on molecular standards (Figure 2A) (see Methods).

**Figure 1.**
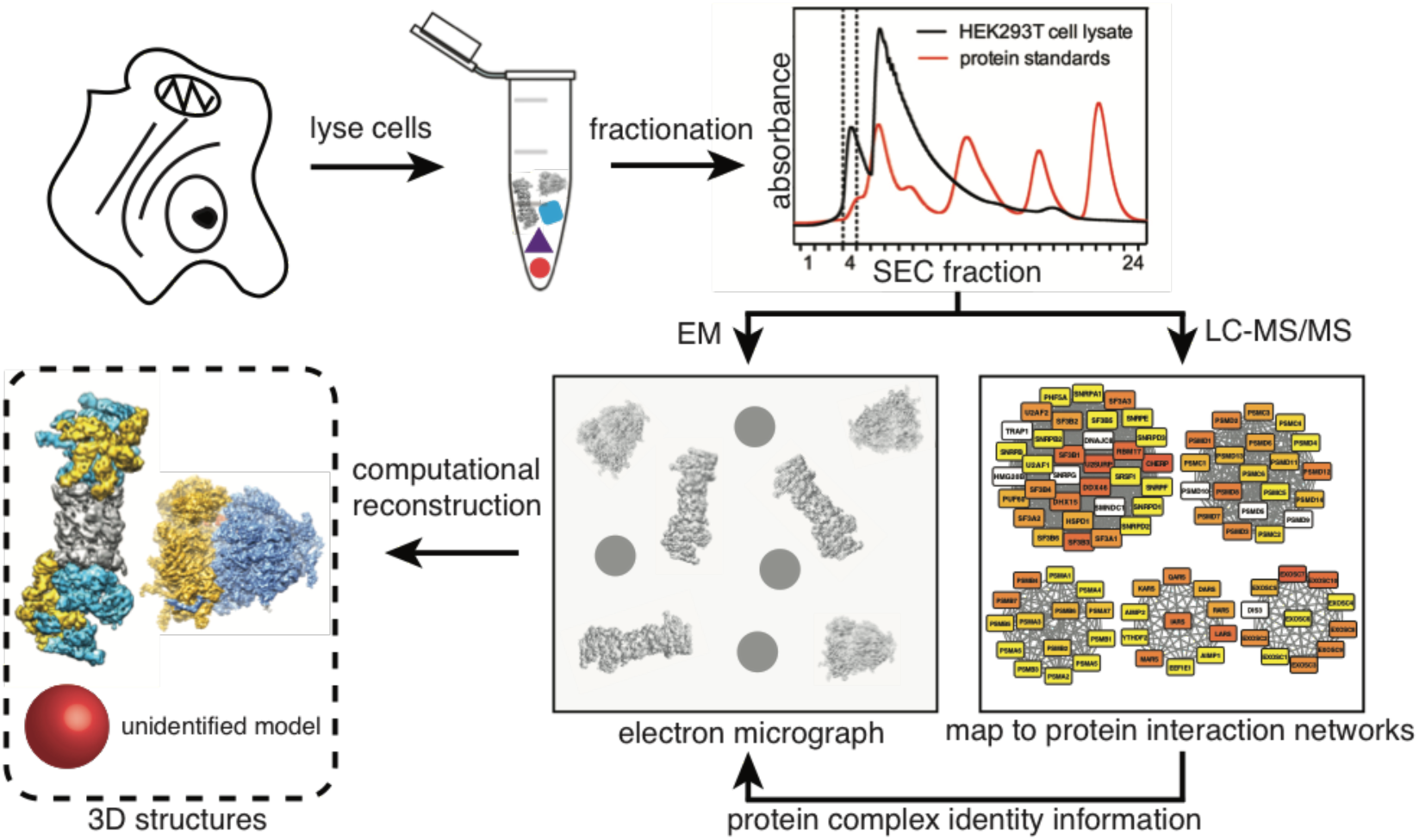
Shotgun EM pipeline used for structural determination of multiple macromolecular complexes. HEK293T cells are subjected to lysis and separation using SEC-FPLC. The resulting fractions are characterized separately by electron microscopy and mass spectrometry. Proteins identified from mass spectrometry are mapped to known and predicted protein complexes to identify which complexes are present in a given fraction. Electron microscopy data is then used to generate structures of multiple protein complexes. Proteasome (EMD-1992) and Ribosome (EMD-2938) shown as examples structures.

**Figure 2.**
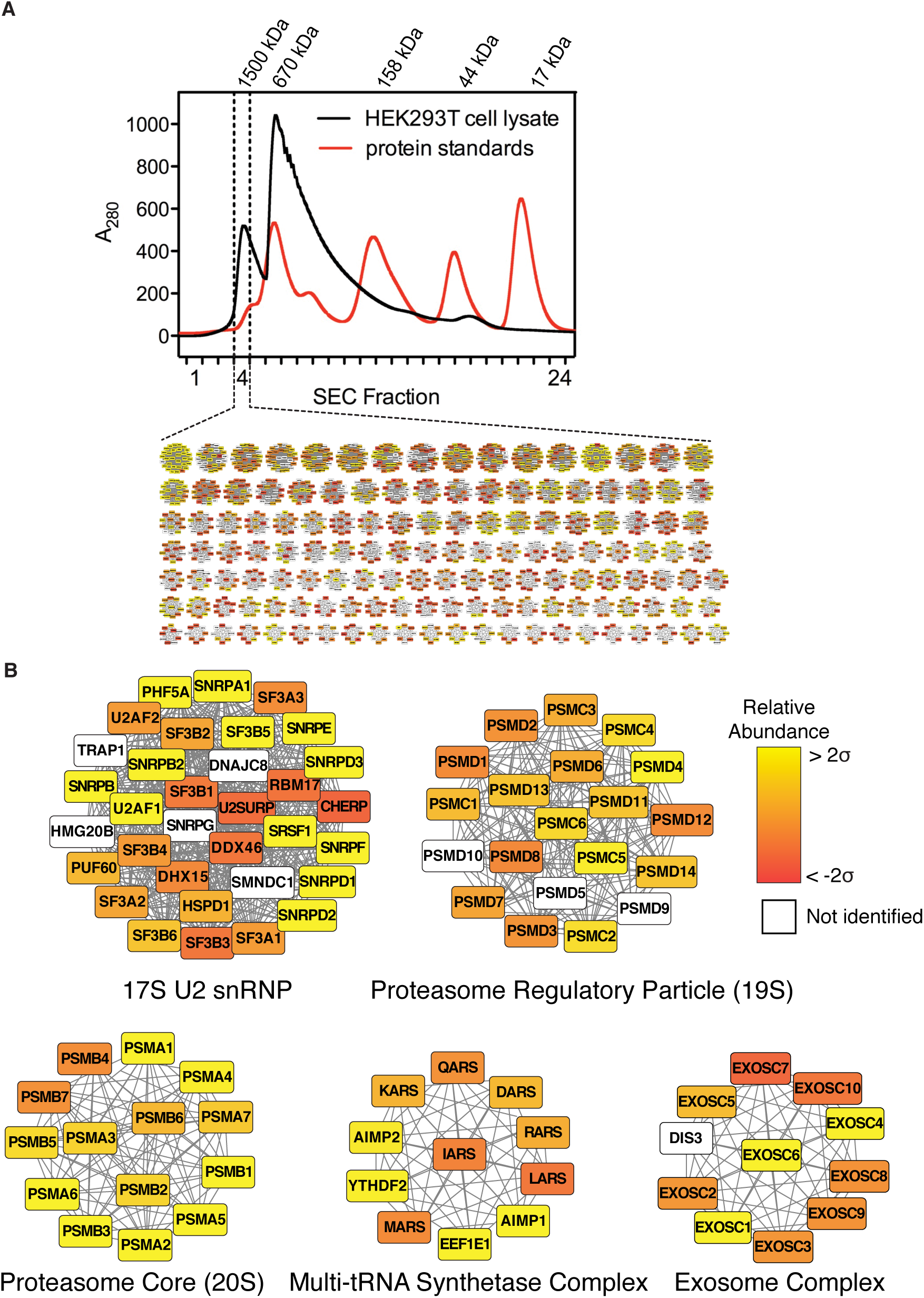
Identification of protein complexes in a cellular fraction. (A) Elution profile from SEC-FPLC. Elution profiles of protein standards are overlaid to estimate the molecular weight range of protein complexes in fraction 4. Inset, a network map displaying a portion of the 1090 candidate complexes determined by mapping mass spectrometry data to CORUM (Ruepp et al., 2010) and hu.MAP (Drew et al., 2017a). (B) Enlarged view of a subset of candidate complexes. A filled node indicates a protein was identified by mass spectrometry, a white node indicates the protein was not identified. Color gradation of filled nodes indicates the relative abundance (determined by label-free quantification of peptide spectral matches) ranging from ±2 standard deviations.

MS analysis of our sample (Figure 2A) identified 1,907 unique proteins. Over 93% of the identified proteins had a molecular weight under 200 kDa indicating that the proteins are likely multi-subunit complexes in order to elute in the high molecular weight fraction. We then mapped proteins identified by MS to a protein-protein interaction network containing previously identified complexes from hu.MAP (Drew et al., 2017a) and CORUM (Ruepp et al., 2010) (Figure 2A). These networks provide a list of documented and high confidence human protein complexes. We identified specific, well annotated protein complexes within our sample, which contains both structurally defined complexes (e.g. the proteasome) (Lander et al., 2012; Schweitzer et al., 2016) and complexes without known structures (e.g. the multi-tRNA synthetase complex) (Mirande, 2017) (Figure 2B).

Complexes with at least 50% of their subunits identified were kept as candidates for subsequent analysis. The resulting list contained 1,090 candidate complexes, many of which share a number of individual subunits and are different variants of the same complex. To identify the most likely complex variant present in our sample, we created a hierarchical network of complexes by performing an all-by-all comparison of proteins between each complex. The resulting network shows related groups of complexes where at least 90% of subunits in higher-order complexes are shared between sub-complexes (Figure S1). Using a threshold of 90% similarity (see Methods), 706 of the 1090 complexes identified by MS belong to a group of shared complexes. Furthermore, the 706 shared complexes in our sample can be organized into 145 distinct hierarchies. This suggests we have 145 groups of related complexes (i.e. with shared subunits) in addition to the remaining 384 unique complexes for a total of 529 complexes in our sample (Table S1).

We then calculated the relative abundance of each complex using label-free quantification (Zybailov et al., 2006) (see Methods). By combining our hierarchical network with the relative abundance for each complex, we were able to identify the specific subunit composition of complexes most likely to be present in our sample. As an example, we can examine the group of related proteasome complexes (Figure S1), showing many related proteasome complexes, where the canonical 20S core appears to be the most abundant form. We then sorted this list of complexes in our sample by relative abundance as a tool to identify which macromolecular machines might be visible by EM.

### Electron microscopy of single particles from HEK293T cell extract fraction

Having identified candidate complexes by MS, we investigated our sample by negative stain EM. Negative stain EM samples are easily prepared and are often used to determine the heterogeneity of a sample. Raw micrographs of our sample show monodisperse particles with clear structural features (Figure 3A). Intact, structurally heterogeneous complexes can be directly observed. The proteasome can be seen in three different structural states, as a core (20S), as a single-capped proteasome (20S core with one 19S regulatory particle) and as a double-capped proteasome (26S, 20S core with two 19S regulatory particles). In addition, many other unidentified particles can be clearly seen, with an average particle size of ~200 Å.

**Figure 3.**
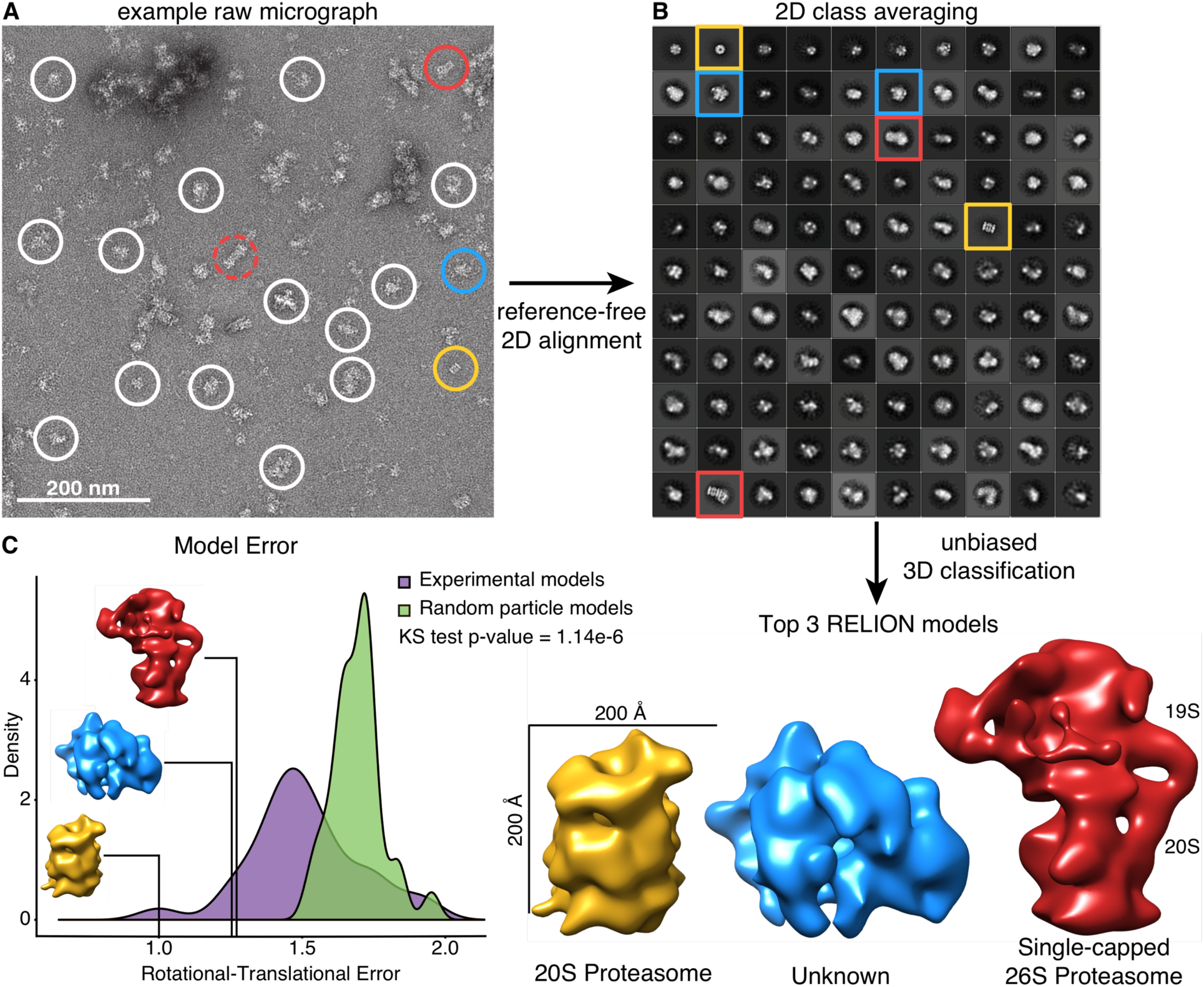
Structural characterization of protein complexes in a cellular fraction. (A) Raw micrograph of negatively-stained sample from SEC-FPLC. Proteasome particles in three different biochemical forms, 20S core, single-capped 26S (20S core with one 19S regulatory particle) and double-capped 26S (20S core with two 19S regulatory particle) circled in gold, red and dashed-red respectively. Representative unidentified particles circled in white. (B) Reference-free 2D class averages of 31,731 template picked particles generated using RELION. The size of each box is 576 Å × 576 Å. The 2D class averages are sorted in decreasing order based on the number of particles belonging to a class with 110 out of 300 2D classes shown. (C) Top 3 models generated using RELION. Models were scored based on their rotational-translational error. The distribution of model error scores was compared to models generated using random particles from our template picked data.

Template picking from 1,250 micrographs of our sample resulted in a final set of 31,731 particles after filtering out ~67% of particles as junk particles (see Methods). To assess the quality of automated template picking, we additionally manually selected 35,381 particles for alignment and classification. A comparison of the reference-free 2D class averages of both manually and template picked data sets yielded similar results (Figure S2), and both data sets were used for independent downstream processing. 2D class averages yielded distinct class averages with various morphologies and features. Remarkably, many well-defined classes emerged from this heterogeneous mixture of complexes (Figure 3B).

Given the success of 2D classification at separating particles into distinct classes, we then proceeded to 3D structure determination using RELION (Scheres, 2012) to generate reconstructions of 30 classes. RELION was developed to group 2D projections of conformationally variable particles into internally consistent groups. Here, we asked if RELION could also classify projections from distinct complexes in a heterogeneous mixture, essentially using RELION to cluster the 2D particles into internally consistent (low error) reconstructions. The 30 3D reconstructions generated all contained various degrees of structural details ranging from distinct barrel shapes to more globular shapes (Figure S2).

To test the internal consistency of 3D reconstructions we determined the distribution of calculated error within the models and ranked each reconstruction based on a rotational-translational error score (see Methods). The error score distribution was then compared to the rotational-translational error scores of models built from random particles in the data set to evaluate our ability to classify related particles belonging to a particular model and demonstrates our 3D reconstructions have substantially less error than random reconstructions (Figure 3C).

We then performed cross-correlations between our top 3 models and several complexes with known structure from our list of high-abundance complexes to determine if we could link our structural models with complex identity (Figure S3) (see Methods). The 20S proteasome emerges as a clear match when compared to our highest scoring model with a cross-correlation score of 0.87. We were also able to distinguish a single-capped proteasome which matched to our third highest scoring model with a cross-correlation score of 0.81. Interestingly, our second highest scoring model was not readily recognizable and none of the known structures emerged as a clear match after cross-correlation. Based on the high-abundance 2D class averages and large volume of the unknown complex, we filtered our proteomics data to search for possible identities. Our search suggests the unknown complex is likely a variant of a mitochondrial ribosome, spliceosome or DNA-repair complex, but given the current resolution, the results are inconclusive. These experiments suggest it is possible to solve multiple structures from cell lysate in a parallel manner, even in the absence of matching starting models.

### Quantification and *ab initio* reconstruction of the proteasome

To determine our ability to further characterize complexes identified in a complex mixture, we investigated our sample specifically in the context of the proteasome, which allowed us to evaluate the success of reconstructions without an initial model. Our goals were to: (1) investigate if *ab initio* reconstructions would reveal clear proteasome structures, (2) determine the ratio of the 20S core and single-capped proteasomes using our single particle data, and (3) compare single particle counting of the proteasome to label-free MS quantification.

Class averages of the 20S core and single-capped proteasomes were clearly identified as barrel-shaped particles and barrels with large rectangular caps, respectively (Figure 4A). Based on identifying the proteasome with notably distinct 2D class averages, as well as RELION-based 3D classification producing two identifiable proteasome models, we asked if *ab initio* reconstructions were capable of correctly recovering proteasome structures. We therefore attempted a completely unsupervised approach for 3D classification using cryoSPARC (Punjani et al., 2017). cryoSPARC was developed for determining multiple 3D structures of a protein without prior structural knowledge or the assumption that the ensemble of conformations resembled each other, but in this context, we evaluated its ability to classify 2D particles of distinct complexes in a mixture. Remarkably, a 3D reconstruction of the 20S core was generated using *ab initio* reconstruction in cryoSPARC with either 5, 10, or 15 classes (Figure S4).

**Figure 4.**
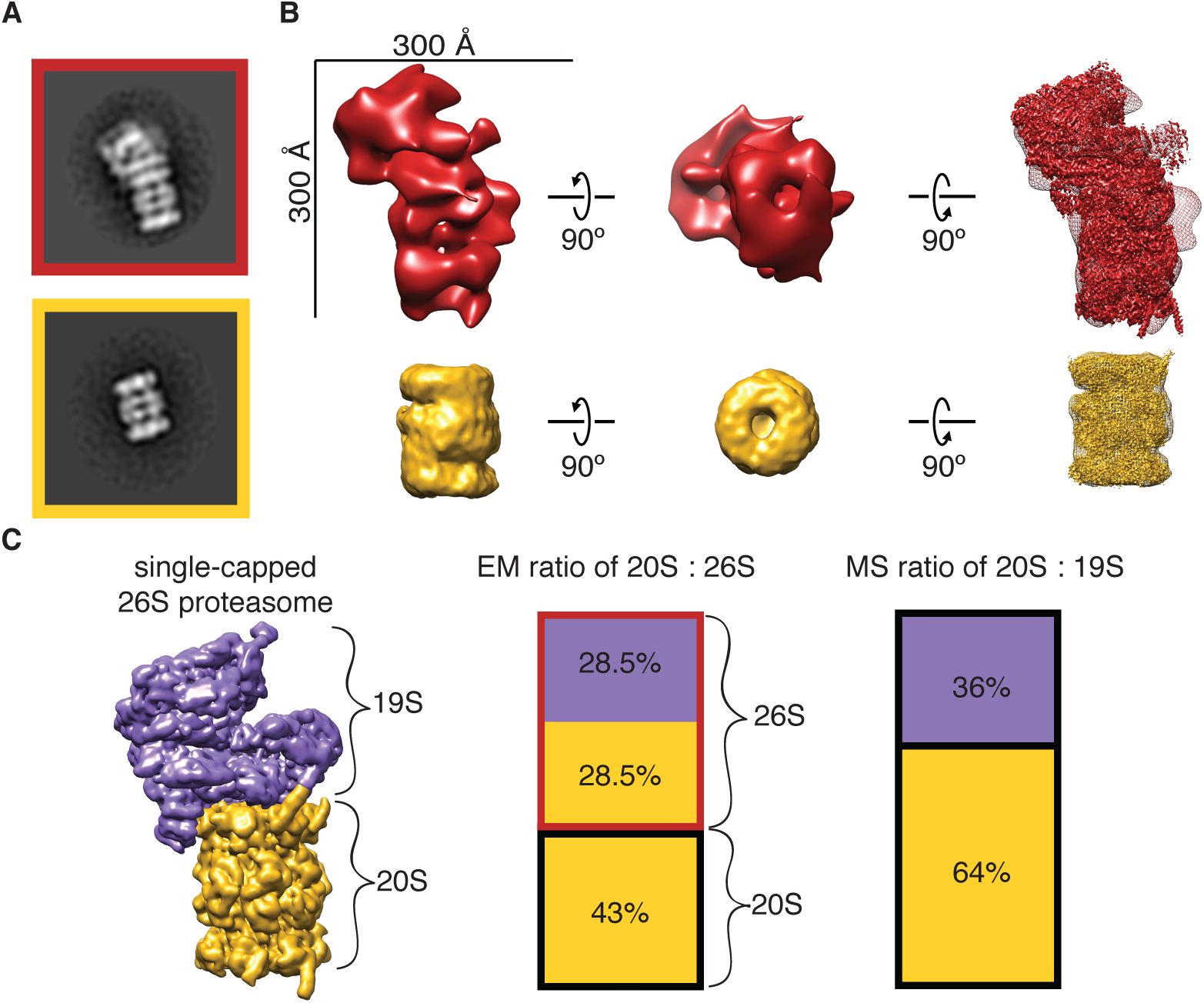
*Ab initio* structures from a cellular fraction unambiguously reveal the proteasome. (A) Reference-free 2D class averages of the proteasome from Figure 3B. (B) Top, structure of single capped proteasome generated using RELION from manually picked particles. Bottom, *ab initio* structure of the 20S core proteasome generated using cryoSPARC. High-resolution structures EMD-4002 (Schweitzer et al., 2016) and EMD-2981 (da Fonseca and Morris, 2015) are fit into the structures, respectively. (C) Quantification of proteasome particles by EM and MS. EMD-1992 (Lander et al., 2012) of 26S proteasome shown as reference.

From the structures generated with 10 classes, a distinct 3D reconstruction of the 20S core showing a clear barrel with a central channel, and some separation of co-axial rings was produced (Figure 4B). This 20S core reconstruction contains 3,150 particles with an estimated resolution of 20.4 Å using the 0.143 Fourier shell correlation (FSC) criterion (Figure S4). Our 3D map is consistent with a recent high-resolution structure of the 20S core (EMD-2981) (da Fonseca and Morris, 2015) with a cross-correlation score of 0.94.

We were unable to distinguish a 3D structure of the single-capped proteasome from cryoSPARC. However, going back to our single-capped proteasome from 3D classification using RELION, we were able to dock in a high-resolution structure determined previously (EMD-4002) (Schweitzer et al., 2016) (Figure 4B). The high-resolution structure can be unambiguously docked into our EM density (cross-correlation score of 0.76) albeit with less agreement given the low number of particles in the model (1,121 particles). Using RELION to refine the structure of our single-capped proteasome, we achieved a nominal resolution of 31 Å (Figure S4).

We then quantified the ratio of 20S core to single-capped proteasome particles by directly counting individual particles from our EM data of fractionated cell lysate. Revisiting our 2D classification, we compared the number of particles aligned in the side view of the 20S core and single-capped proteasome (Figure 4A). The ratio of 20S core to single-capped proteasome particles in our sample was calculated to be 3:2, or 1 bound 19S regulatory particles for every 2.5 20S core particles in our sample by EM. This is consistent with our MS data which suggests the ratio of 19S regulatory particles to 20S core particles is 4:7 (approximately one regulatory particle for every two cores) (Figure 4C). Collectively, our study suggests it is not only possible to solve structures of protein complexes from cell lysate *ab initio*, but also quantify the stoichiometry of biochemical states.

## Discussion

One bottleneck of structural biology is studying only a single protein or protein complex structure in a single experiment. However, recent advances in detectors and software for EM brings about the possibility of high-throughput structural determination using EM. To this end, we have demonstrated shotgun EM as a potential pipeline for high-throughput identification and structural determination of macromolecular machines. By combining MS and EM, we demonstrate it is possible to structurally characterize and identify protein complexes from a cellular sample containing many native complexes. This pipeline was used to successfully identify the proteasome from a cellular fraction with minimal user input. Additionally, we construct a self-consistent structural model of an ~1 MDa protein complex of unknown identity.

A recent study showed that higher-order assemblies from a eukaryotic thermophile could be separated chromatographically, identified by MS and visualized through cryo-electron microscopy to obtain a high resolution structure (Kastritis et al., 2017). The authors performed cryo-EM on particles from a complex mixture to solve a 4.7 Å resolution structure of fatty acid synthase from cell lysate separated by molecular size after a 50% enrichment for fatty acid synthase. In our study using human cells, which have a canonical proteome approximately 3 times larger than *C. thermophilum*, we are able to obtain structural information from a complex mixture, suggesting that sample heterogeneity is a surmountable problem.

By classifying heterogeneous particles from a cellular fraction, we show that structural studies on heterogeneous mixtures are possible but several key barriers remain, with the main challenge being to correctly assign different orientations of the same complex in large data sets of heterogeneous mixtures. While currently we cannot identify each class average or 3D structure obtained in this study, we are able to distinguish different structural states of the proteasome using current *ab initio* methods suggesting that shotgun EM is a promising tool to characterize the heterogeneity of protein complex forms. Our unidentified top scoring model was not readily recognizable and had no apparent match from model fitting. It is possible our model has been structurally annotated previously but was not covered in our search. Alternatively, it is possible our model remains unidentified because it is structurally novel. In future experiments, a comprehensive list of solved structures coupled with optimal volume alignment and cross-correlation can be used to identify likely matches to models generated using shotgun EM.

One challenge when dealing with protein complexes is defining their precise subunits. MS does not indicate which complex a protein belonging to multiple complexes was identified from. Many of these related complexes and sub-complexes have yet to be structurally or biochemically characterized. Our hierarchical network strategy allows us to make an initial estimate on which form of a complex might be in our EM data. Using shotgun EM, we aim to validate these uncharacterized and other less characterized forms of complexes that may be more amenable to our separation scheme.

A key proof of concept in this study was the proteasome, which is a structurally distinct complex and serves a crucial role in protein degradation in eukaryotic cells (Finley, 2009). The native stoichiometry of the proteasome has been studied in different ways by multiple groups (Asano et al., 2015; Havugimana et al., 2012). Our template-picked counting of single proteasome particles has an advantage over MS approaches by identifying which form of a complex an identified protein belongs to. While our MS and EM quantification were consistent showing an approximate ratio of 20S core to 19S regulatory particles of 2:1, a separate study using corrected spectral counts suggests the ratio is closer to 4:1 (Havugimana et al., 2012). To reconcile these two observations, more chromatographic fractions containing the proteasome would need to be quantified by EM and MS to see if there is agreement. As more protein complexes become structurally annotated, shotgun EM can be used as an auxiliary method for quantifying the abundances of native complexes, as well as their stoichiometry.

After *ab initio* 3D classification, we obtained a reasonable reconstruction of the 20S core in cryoSPARC from 3,150 particles. Although only half of these particles are accounted for from 2D class averaging of all particles, it is likely that the discrepancy results from proteasome particles that are misclassified or exist in different, less-populated orientations in our 2D class averages. Alternatively, because the number of models we could reconstruct in 3D was limited by the small populations of each complex we had in our micrographs, it is possible that non-proteasome particles were grouped into our 3D class of the proteasome. These misclassified particles would have a small contribution to the overall likelihood of the 3D map as it is reconstructed (Punjani et al., 2017). One method to separate misclassified particles would be to do iterative rounds of 3D classification.

In this study, we used a 60S ribosome class average as a template for auto-picking due to its large molecular weight and round shape. Interestingly, none of the resulting averages resembled the 60S, providing evidence that we were not biasing the results from template picking and subsequent data analysis. A similar concern for model bias exists when using RELION to generate 3D models. Despite this, none of the 3D classes are visually identical to the reference 3D model, with most EMD structures selected from our MS data outscoring the reference model by cross-correlation score when compared to our top 3 RELION models. In future experiments, more sophisticated template matching, deep learning algorithms, or *ab initio* methods can be introduced to improve particle identification and model building (Punjani et al., 2017; Rickgauer et al., 2017; Wang et al., 2016).

This study represents a foray into structural proteomics using electron microscopy, suggesting that parallel structural determination of protein complexes shows promise for alleviating bottlenecks in structural biology. In the interim before high-resolution data is collected, it is possible to search for structurally uncharacterized complexes through the addition of protein tags (Flemming et al., 2010) to identify complexes in a heterogeneous mix without the need to purify the sample. One could also utilize integrative structural biology approaches to have a predicted model with which to search for structures in cell extract. We envision using cryo-electron microscopy for this pipeline to solve sub-nanometer resolution structures, where homology models and known structures can be more clearly compared.

Shotgun EM will accelerate the pace at which structural information is generated, and allow us to better understand the structure-function relationship of proteins. Optimization of this technique has the potential to address questions about many macromolecular machines across different cell types, disease states, and species. We propose that investigating the collective protein complexes in a cell, or the ‘complexome’, using shotgun cryo-EM will help inform us broadly on systems biology, cell biology and changes in complexes that contribute to human diseases.

## Author Contributions

E.J.V. performed electron microscopy, single particle classification and reconstruction. K.D. and E.J.V performed bioinformatics analysis. A.L.M performed cellular fractionation and mass spectrometry. E.J.V., A.L.M., K.D., E.M.M., and D.W.T. analyzed and interpreted the data and wrote the manuscript. D.W.T., E.M.M, A.L.M, K.D., and E.J.V. conceived of experiments. E.M.M. and D.W.T. supervised the study and secured funding for the work.

## Acknowledgements

We thank A. Johnson for kindly providing 60S ribosomes used to create templates; A. Matouschek for critical reading of the manuscript; and members of the Marcotte and Taylor labs for helpful discussions. MS was acquired in the UT Austin Proteomics Facility supported by CPRIT RP110782 to Maria Person. This work was supported in part by Welch Foundation Grants F-1938 (to D.W.T.) and F-1515 (to E.M.M), NIH grants F32 GM112495, K99 HD092613 (to K.D.) and R35 GM122480, DP1 GM106408, R21 GM119021, R01 DK110520, R01 HD085901 (to E.M.M), NSF IOS-1237975 (to E.M.M), and Army Research Office W911NF-12-1-0390 (to E.M.M). D.W.T is a CPRIT Scholar supported by the Cancer Prevention and Research Institute of Texas (RR160088).

## References

Asano, S., Fukuda, Y., Beck, F., Aufderheide, A., Förster, F., Danev, R., and Baumeister, W. (2015). A molecular census of 26S proteasomes in intact neurons. Science 347, 439–442.

Chandonia, J.-M., and Brenner, S.E. (2006). The impact of structural genomics: expectations and outcomes. Science 311, 347–351.

Danev, R., and Baumeister, W. (2016). Cryo-EM single particle analysis with the Volta phase plate. Elife 5, e13046.

Drew, K., Lee, C., Huizar, R.L., Tu, F., Borgeson, B., McWhite, C.D., Ma, Y., Wallingford, J.B., and Marcotte, E.M. (2017a). Integration of over 9,000 mass spectrometry experiments builds a global map of human protein complexes. Mol. Syst. Biol. 13, 932.

Drew, K., Müller, C.L., Bonneau, R., and Marcotte, E.M. (2017b). Identifying direct contacts between protein complex subunits from their conditional dependence in proteomics datasets. PLOS Comput. Biol. 13, e1005625.

Finley, D. (2009). Recognition and Processing of Ubiquitin-Protein Conjugates by the Proteasome. Annu. Rev. Biochem. 78, 477–513.

Flemming, D., Thierbach, K., Stelter, P., Böttcher, B., and Hurt, E. (2010). Precise mapping of subunits in multiprotein complexes by a versatile electron microscopy label. Nat. Struct. Mol. Biol. 17, 775–778.

da Fonseca, P.C.A., and Morris, E.P. (2015). Cryo-EM reveals the conformation of a substrate analogue in the human 20S proteasome core. Nat. Commun. 6, 7573.

Gavin, A.-C., Bösche, M., Krause, R., Grandi, P., Marzioch, M., Bauer, A., Schultz, J., Rick, J.M., Michon, A.-M., and Cruciat, C.-M. (2002). Functional organization of the yeast proteome by systematic analysis of protein complexes. Nature 415, 141–147.

Havugimana, P.C., Hart, G.T., Nepusz, T., Yang, H., Turinsky, A.L., Li, Z., Wang, P.I., Boutz, D.R., Fong, V., Phanse, S., et al. (2012). A Census of Human Soluble Protein Complexes. Cell 150, 1068–1081.

Hein, M.Y., Hubner, N.C., Poser, I., Cox, J., Nagaraj, N., Toyoda, Y., Gak, I.A., Weisswange, I., Mansfeld, J., Buchholz, F., et al. (2015). A Human Interactome in Three Quantitative Dimensions Organized by Stoichiometries and Abundances. Cell 163, 712–723.

Ho, Y., Gruhler, A., Heilbut, A., Bader, G.D., Moore, L., Adams, S.-L., Millar, A., Taylor, P., Bennett, K., Boutilier, K., et al. (2002). Systematic identification of protein complexes in Saccharomyces cerevisiae by mass spectrometry. Nature 415, 180–183.

Huttlin, E.L., Ting, L., Bruckner, R.J., Gebreab, F., Gygi, M.P., Szpyt, J., Tam, S., Zarraga, G., Colby, G., Baltier, K., et al. (2015). The BioPlex Network: A Systematic Exploration of the Human Interactome. Cell 162, 425–440.

Huttlin, E.L., Bruckner, R.J., Paulo, J.A., Cannon, J.R., Ting, L., Baltier, K., Colby, G., Gebreab, F., Gygi, M.P., Parzen, H., et al. (2017). Architecture of the human interactome defines protein communities and disease networks. Nature 545, 505–509.

Kastritis, P.L., O’Reilly, F.J., Bock, T., Li, Y., Rogon, M.Z., Buczak, K., Romanov, N., Betts, M.J., Bui, K.H., Hagen, W.J., et al. (2017). Capturing protein communities by structural proteomics in a thermophilic eukaryote. Mol. Syst. Biol. 13, 936.

Kristensen, A.R., Gsponer, J., and Foster, L.J. (2012). A high-throughput approach for measuring temporal changes in the interactome. Nat. Methods 9, 907–909.

Krogan, N.J., Cagney, G., Yu, H., Zhong, G., Guo, X., Ignatchenko, A., Li, J., Pu, S., Datta, N., Tikuisis, A.P., et al. (2006). Global landscape of protein complexes in the yeast Saccharomyces cerevisiae. Nature 440, 637–643.

Kühlbrandt, W. (2014). The resolution revolution. Science 343, 1443–1444.

Lander, G.C., Stagg, S.M., Voss, N.R., Cheng, A., Fellmann, D., Pulokas, J., Yoshioka, C., Irving, C., Mulder, A., Lau, P.-W., et al. (2009). Appion: An integrated, database-driven pipeline to facilitate EM image processing. J. Struct. Biol. 166, 95–102.

Lander, G.C., Estrin, E., Matyskiela, M.E., Bashore, C., Nogales, E., and Martin, A. (2012). Complete subunit architecture of the proteasome regulatory particle. Nature.

Leitner, A., Faini, M., Stengel, F., and Aebersold, R. (2016). Crosslinking and Mass Spectrometry: An Integrated Technology to Understand the Structure and Function of Molecular Machines. Trends Biochem. Sci. 41, 20–32.

Liu, F., and Heck, A.J. (2015). Interrogating the architecture of protein assemblies and protein interaction networks by cross-linking mass spectrometry. Curr. Opin. Struct. Biol. 35, 100–108.

Mastronarde, D.N. (2005). Automated electron microscope tomography using robust prediction of specimen movements. J. Struct. Biol. 152, 36–51.

Mirande, M. (2017). The Aminoacyl-tRNA Synthetase Complex. In Macromolecular Protein Complexes, J.R. Harris, and J. Marles-Wright, eds. (Cham: Springer International Publishing), pp. 505–522.

Montelione, G.T. (2012). The Protein Structure Initiative: achievements and visions for the future. F1000 Biol. Rep. 4.

Pettersen, E.F., Goddard, T.D., Huang, C.C., Couch, G.S., Greenblatt, D.M., Meng, E.C., and Ferrin, T.E. (2004). UCSF Chimera—A visualization system for exploratory research and analysis. J. Comput. Chem. 25, 1605–1612.

Punjani, A., Rubinstein, J.L., Fleet, D.J., and Brubaker, M.A. (2017). cryoSPARC: algorithms for rapid unsupervised cryo-EM structure determination. Nat. Methods 14, 290–296.

Rappsilber, J., Siniossoglou, S., Hurt, E.C., and Mann, M. (2000). A Generic Strategy To Analyze the Spatial Organization of Multi-Protein Complexes by Cross-Linking and Mass Spectrometry. Anal. Chem. 72, 267–275.

Rickgauer, J.P., Grigorieff, N., and Denk, W. (2017). Single-protein detection in crowded molecular environments in cryo-EM images. ELife 6.

Rohou, A., and Grigorieff, N. (2015). CTFFIND4: Fast and accurate defocus estimation from electron micrographs. J. Struct. Biol. 192, 216–221.

Roseman, A. (2004). FindEM—a fast, efficient program for automatic selection of particles from electron micrographs. J. Struct. Biol. 145, 91–99.

Ruepp, A., Waegele, B., Lechner, M., Brauner, B., Dunger-Kaltenbach, I., Fobo, G., Frishman, G., Montrone, C., and Mewes, H.-W. (2010). CORUM: the comprehensive resource of mammalian protein complexes—2009. Nucleic Acids Res. 38, D497–D501.

Scheres, S.H.W. (2012). RELION: Implementation of a Bayesian approach to cryo-EM structure determination. J. Struct. Biol. 180, 519–530.

Schweitzer, A., Aufderheide, A., Rudack, T., Beck, F., Pfeifer, G., Plitzko, J.M., Sakata, E., Schulten, K., Förster, F., and Baumeister, W. (2016). Structure of the human 26S proteasome at a resolution of 3.9 Å. Proc. Natl. Acad. Sci. 113, 7816–7821.

Shannon, P., Markiel, A., Ozier, O., Baliga, N.S., Wang, J.T., Ramage, D., Amin, N., Schwikowski, B., and Ideker, T. (2003). Cytoscape: a software environment for integrated models of biomolecular interaction networks. Genome Res. 13, 2498–2504.

Sibanda, B.L., Chirgadze, D.Y., Ascher, D.B., and Blundell, T.L. (2017). DNA-PKcs structure suggests an allosteric mechanism modulating DNA double-strand break repair. Science 355, 520–524.

Suloway, C., Pulokas, J., Fellmann, D., Cheng, A., Guerra, F., Quispe, J., Stagg, S., Potter, C.S., and Carragher, B. (2005). Automated molecular microscopy: The new Leginon system. J. Struct. Biol. 151, 41–60.

Vaudel, M., Barsnes, H., Berven, F.S., Sickmann, A., and Martens, L. (2011). SearchGUI: An open-source graphical user interface for simultaneous OMSSA and X!Tandem searches. PROTEOMICS 11, 996–999.

Vaudel, M., Burkhart, J.M., Zahedi, R.P., Oveland, E., Berven, F.S., Sickmann, A., Martens, L., and Barsnes, H. (2015). PeptideShaker enables reanalysis of MS-derived proteomics data sets. Nat. Biotechnol. 33, 22–24.

Wan, C., Borgeson, B., Phanse, S., Tu, F., Drew, K., Clark, G., Xiong, X., Kagan, O., Kwan, J., Bezginov, A., et al. (2015). Panorama of ancient metazoan macromolecular complexes. Nature 525, 339–344.

Wang, F., Gong, H., Liu, G., Li, M., Yan, C., Xia, T., Li, X., and Zeng, J. (2016). DeepPicker: A deep learning approach for fully automated particle picking in cryo-EM. J. Struct. Biol. 195, 325–336.

Zybailov, B., Mosley, A.L., Sardiu, M.E., Coleman, M.K., Florens, L., and Washburn, M.P. (2006). Statistical Analysis of Membrane Proteome Expression Changes in Saccharomyces cerevisiae. J. Proteome Res. 5, 2339–2347.

